# Adaptations in common synaptic inputs to spinal motor neurons during grasping versus a less functional hand task

**DOI:** 10.1101/2025.05.02.651931

**Authors:** Hélio V. Cabral, Caterina Cosentino, Andrea Rizzardi, J. Greig Inglis, Andrew J. Fuglevand, Francesco Negro

## Abstract

Previous evidence suggests that shared synaptic inputs across spinal motor neurons play a key role in coordinating multiple muscles during hand movements, reducing control complexity. In this study, we investigated how the nervous system modulates these common synaptic inputs during a functionally relevant grip (grasping) compared to less functionally relevant hand tasks. Seventeen participants performed three different tasks: simultaneous four-finger flexion without thumb involvement (four-finger flexion), thumb flexion, and simultaneous flexion of both fingers and thumb (grasping). For each task, subjects sustained isometric contractions at 5% and 15% of maximal voluntary contraction, while high-density surface electromyograms (HDsEMG) were recorded from the superficial extrinsic flexor muscles of the hand. Motor unit spike trains were decomposed from HDsEMG and tracked across tasks, and their mean discharge rate was calculated. Coherence between motor units was quantified within the delta, alpha, and beta bands to estimate common synaptic oscillations. At both force levels, the mean discharge rate decreased during grasping compared to four-finger flexion but increased during grasping compared to thumb flexion. Additionally, the area under the curve of coherence within the alpha band decreased by ∼20% during grasping compared to the four-finger flexion task, with no significant delta or beta bands changes. These reductions in alpha band coherence were reflected in force oscillations, showing decreased force-neural drive coupling within the alpha band and increased force steadiness during grasping compared to four-finger flexion. Our findings suggest that a functionally relevant and frequently used grip involves distinct neural control mechanisms that ultimately enhance force control.

## Introduction

The human hand is capable of performing a wide range of movements with remarkable versatility, enabling complex interactions with the environment (Napier, 1956). While humans possess some degree of finger individualization, most hand movements rely on the coordinated flexion of multiple fingers, particularly for grasping, a fundamental action in daily activities (Young, 2003, Almécija et al., 2015, Schieber and Santello, 2004). The anatomical specialization of the human hand, such as a fully opposable thumb, larger intrinsic muscles, and the ability to arch the palm of the hand, supports the performance of prehensile movements, allowing for efficient object manipulation (Napier, 1956, Napier, 1960, Napier, 1965, Marzke, 1997, Susman, 1979, Marzke and Shackley, 1986). Furthermore, the functional relevance of grasping is evolutionarily evident from early development, as newborns display an instinctive grasping reflex that later transitions into voluntary grasping, reinforcing its role as a fundamental component of human motor behaviour (Dennis, 1943, Twitchell, 1965). Taken together, these findings highlight that while humans exhibit some degree of independent finger control, most everyday manual tasks, from tool use to object manipulation, rely on the coordinated flexion of multiple digits, making grasping one of the most functionally relevant hand movements.

Precision and dexterity during grasping require a finely coordinated interplay of the forces produced by the multiple muscles controlling the fingers (Landsmeer, 1963, Long et al., 1970). Rather than individually controlling each extrinsic hand muscle, the central nervous system synergistically coordinates them, resulting in the required hand movement, which reduces the degrees of freedom and, consequently, simplifies control (for review, see Santello et al. (2013)). This modular control of fingers and hand movements, referred to as “hand synergies”, has been extensively documented through different methodologies, including kinematic data (Soechting and Flanders, 1997, Santello et al., 1998, Santello et al., 2002), force measurements (Li et al., 1998, Reilly and Hammond, 2000, Santello and Soechting, 2000), and muscle activation recordings (Maier and Hepp-Reymond, 1995, Weiss and Flanders, 2004, Tanzarella et al., 2021). These studies demonstrate that a small set of low-dimensional components can account for a wide range of hand movements. This has also been observed at the neuronal level, where synaptic inputs largely shared across spinal motor neurons, particularly in extrinsic hand muscles (Hockensmith et al., 2005, Winges and Santello, 2004), are thought to underlie this modular control (Keen and Fuglevand, 2004, Hockensmith et al., 2005, Winges and Santello, 2004, Johnston et al., 2005, Tanzarella et al., 2021). For example, Hockensmith et al. (2005) reported a high degree of commonality in motor unit discharge times between two extrinsic hand muscles acting on the thumb and index finger. Interestingly, this between-muscle commonality was as pronounced as that observed in motor units within individual muscles. However, it remains less clear how the nervous system modulates these common synaptic inputs across different hand movements, particularly when comparing a frequently used grip, such as grasping, with a less functionally relevant hand action, like the simultaneous flexion of all four fingers without involvement of the thumb.

In this study, we sought to assess the common synaptic oscillations to spinal motor neurons of the extrinsic finger flexor muscles during three tasks: simultaneous flexion of the metacarpophalangeal joints of four fingers without involvement of the thumb (hereafter referred to as four-finger flexion), thumb flexion, and simultaneous flexion of both fingers and thumb (hereafter referred to as grasping). To achieve this, we decomposed motor unit spike trains from high-density surface electromyograms (HDsEMG) recorded from the superficial extrinsic finger flexors and tracked motor units across tasks. We then used coherence analysis to estimate the shared synaptic oscillations in the delta, alpha, and beta bands. Our results revealed a reduction in alpha band oscillations during grasping compared to four-finger flexion. This reduction in alpha band oscillations was reflected in force oscillations, revealing decreased force-neural drive coupling within the alpha band and increased force steadiness during grasping. These findings suggest that a more natural, functionally relevant grip involves specific neural control mechanisms to enhance force control.

## Methods

### Participants

Seventeen healthy individuals (5 females; mean ± standard deviation: age 30 ± 4 years) with no history of upper limb injuries volunteered to participate in this study. Before starting the experiments, all participants provided written informed consent. The experimental procedures received approval from the local ethics committee at the University of Brescia (code NP5665) and were conducted in accordance with the latest version of the Declaration of Helsinki.

### Experimental protocol

The study comprised a single experimental session lasting approximately 1.5 hrs. Participants were seated comfortably in a chair with their right forearm supported on a custom-built device which was secured to the testing table (**Figure 1A**). The elbow was flexed at 45° (0° being the anatomical position), and the wrist was positioned neutrally (i.e., midway between full supination and full pronation). Both the forearm and wrist were secured to the device with Velcro straps. The four fingers were aligned with the forearm (i.e., pointing forward), and the thumb was held in a mid-flexed position. The distal segments of the thumb and fingers were fixed to adjustable supports attached to the load cells (thumb: SM-100 N; fingers: SM-500 N, Interface, Arizona, USA) to record the isometric forces produced by simultaneous flexion of the four fingers and thumb, either separately or simultaneously (**Figure 1A**).

**Figure 1:**
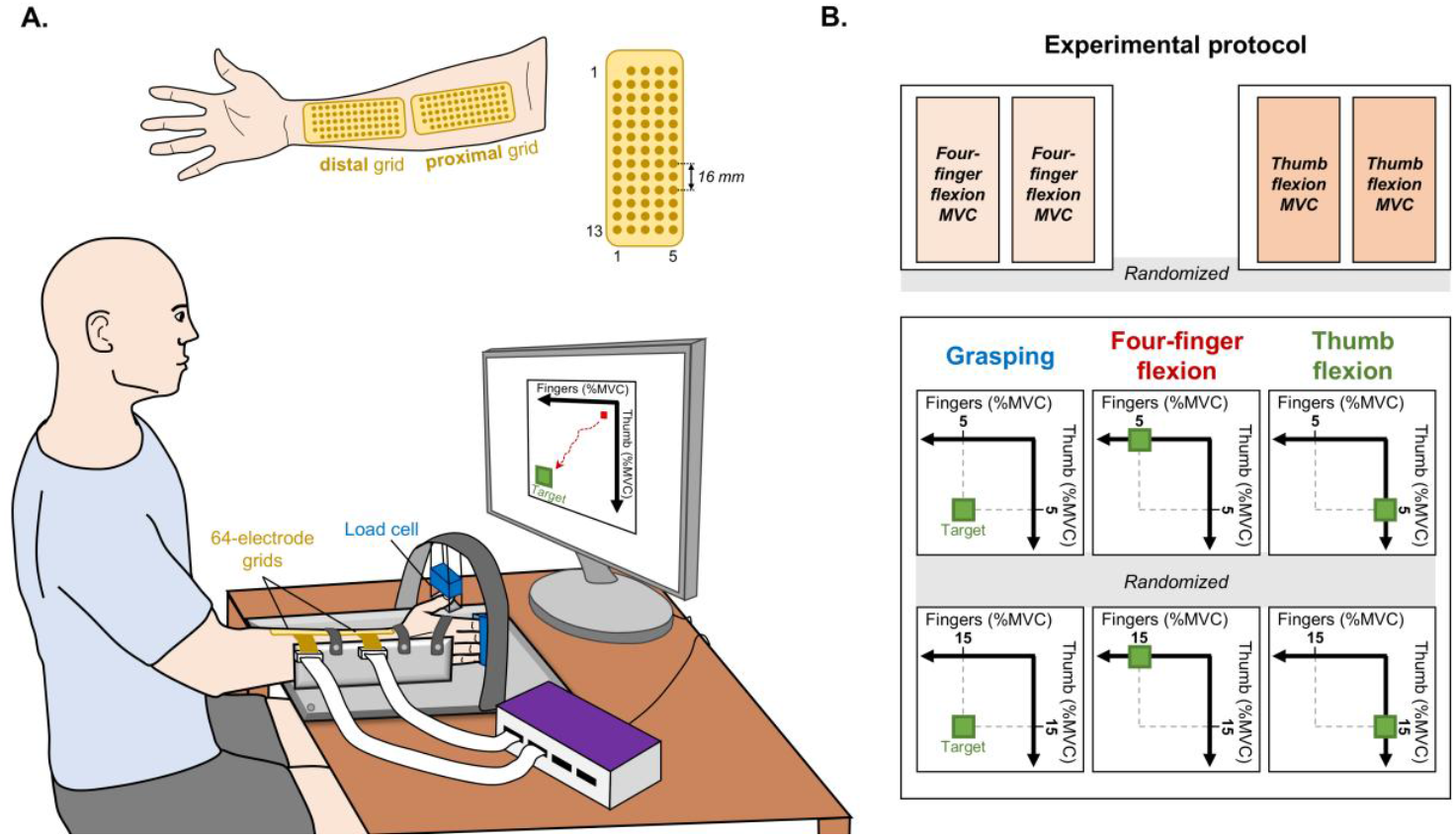
Experimental setup and protocol. A, positioning of participants to perform the experimental task. The distal segments of the fingers and the thumb were fixed to adjustable supports attached to load cells to measure isometric forces produced by simultaneous flexion of the four fingers and thumb, either separately or simultaneously. Two high-density grids of 64 electrodes each were used to record the muscle activity from the superficial extrinsic flexor muscles of the hand (top panel). B, participants performed maximal voluntary isometric contractions (MVC) of four-finger and thumb flexion (top), followed by isometric force-matching contractions (bottom) involving three different tasks: simultaneous flexion of the four fingers without involvement of the thumb (i.e., four-finger flexion), thumb flexion, and simultaneous flexion of both fingers and thumb (i.e., grasping). For each movement, two force levels were performed, 5% and 15% MVC. Visual feedback was provided as a two-dimensional coordinate plane, where the x- and y-axis corresponded to the flexion forces of the four-fingers and thumb, respectively, and a green square indicated the target.

Initially, participants performed maximal voluntary isometric contractions (MVCs) for 3 s, with a 2 min interval of rest in between. Two MVCs of finger flexion and two MVCs of thumb flexion were performed in a randomized order (top panel of **Figure 1B**). For each action, the highest value from the two MVCs was considered the maximal isometric force and used as a reference for subsequent submaximal contractions. Participants were then instructed to perform six isometric force-matching contractions involving three different tasks: simultaneous flexion of the four fingers without involvement of the thumb (i.e., four-finger flexion), thumb flexion, and simultaneous flexion of all fingers with the thumb (i.e., grasping). For each task, participants performed two steady isometric contractions at two target forces: 5% and 15% MVC. Visual feedback of the produced forces was provided as a moving red square in a two-dimensional coordinate plane, where the x- and y-axis corresponded to the forces of four-finger flexion and thumb flexion, respectively, and a green square indicated the target (bottom panel of **Figure 1B**). Participants were required to sustain the force at the target level for 50 s within a tolerance of ± 5%, indicated by the borders of the green square. During the individual movements (four-finger or thumb flexion), the force acquired by the uninvolved load cell was recorded but not displayed on the monitor. The computer monitor displaying the target and produced forces was positioned ∼60 cm in front of the participant.

### Data collection

For all tasks, HDsEMG were recorded in monopolar mode from the superficial extrinsic flexor muscles of the hand using two grids of 64 electrodes each (GR08MM1305; 8 mm inter-electrode distance; OT Bioelettronica, Turin, Italy; **Figure 1A**). To cover as much of the surface area of the muscles as possible, the electrodes were positioned from proximal to distal over the volar surface of the forearm (**Figure 1A**). Before electrode placement, the skin was shaved and mildly abraded (EVERI, Spes Medica, Genova, Italy), and the surface was cleaned with water. The electrode grids were then fixed to the skin using a double-sided adhesive foam, and a conductive paste (AC cream, Spes Medica, Genova, Italy) was used to fill the wells in the foam to ensure optimal electrode-skin contact. A strap electrode dampened with water was used as a reference electrode and positioned on the involved wrist. HDsEMG were sampled at 2048 Hz using a 16-bit amplifier (10-500 Hz bandwidth; Quattrocento, OT Bioelettronica, Turin, Italy). Force output from the load cell was amplified by a variable factor (MISO II, OT Bioelettronica) and sampled synchronously with HDsEMG.

### Data analysis

Data were analyzed offline using MATLAB custom-written scripts (version 2022b, The MathWorks, Natick, MA, USA).

#### Identification and tracking of motor units

The monopolar HDsEMG were bandpass filtered between 20 and 500 Hz using a 3^rd^ order Butterworth filter. The filtered signals were then visually inspected, and channels with artifacts or contact problems were excluded. Subsequently, the HDsEMG from each electrode grid were decomposed separately into motor unit spike trains using a convolutive blind-source separation algorithm (**Figure 2A**) (Negro et al., 2016). This algorithm has been validated previously (Negro et al., 2016) and widely applied in the literature to assess single motor unit activity in upper limb muscles (Kapelner et al., 2018, Maillet et al., 2022, Cabral et al., 2024a). Missing or misidentified motor unit pulses were iteratively edited by an expert operator, followed by the re-estimation of motor unit spike trains (Martinez-Valdes et al., 2017, Hug et al., 2021). After motor unit editing, the motor units were tracked between four-finger flexion and grasping tasks, as well as between thumb flexion and grasping tasks, by reapplying the motor unit separation vectors from one contraction to the other, and vice-versa (Oliveira and Negro, 2021, Rossato et al., 2022, Cabral et al., 2024a). The motor unit separation vectors are distinct for each individual motor unit and serve as spatio-temporal matched filters to estimate motor unit spike trains. Consequently, the same motor units were analyzed across four-finger flexion and grasping tasks, as well as between thumb flexion and grasping tasks. **Figure 2B** shows examples of the innervation pulse trains (i.e., motor unit discharge times) and the two-dimensional representations of the motor unit action potentials of motor units tracked between tasks. For each matched motor unit, the instantaneous discharge rate was obtained as the multiplicative inverse of the interspike intervals, and the average value over the 50-s steady force was calculated (i.e., mean discharge rate) and retained for further analysis. It is important to note that common units from both the proximal and distal electrode grids were identified, and the one with the highest coefficient of variation of the interspike interval was removed from the analysis. Motor unit spike trains were considered to be generated by the same motor unit if they had at least 30% of common discharge times (Negro et al., 2016).

**Figure 2:**
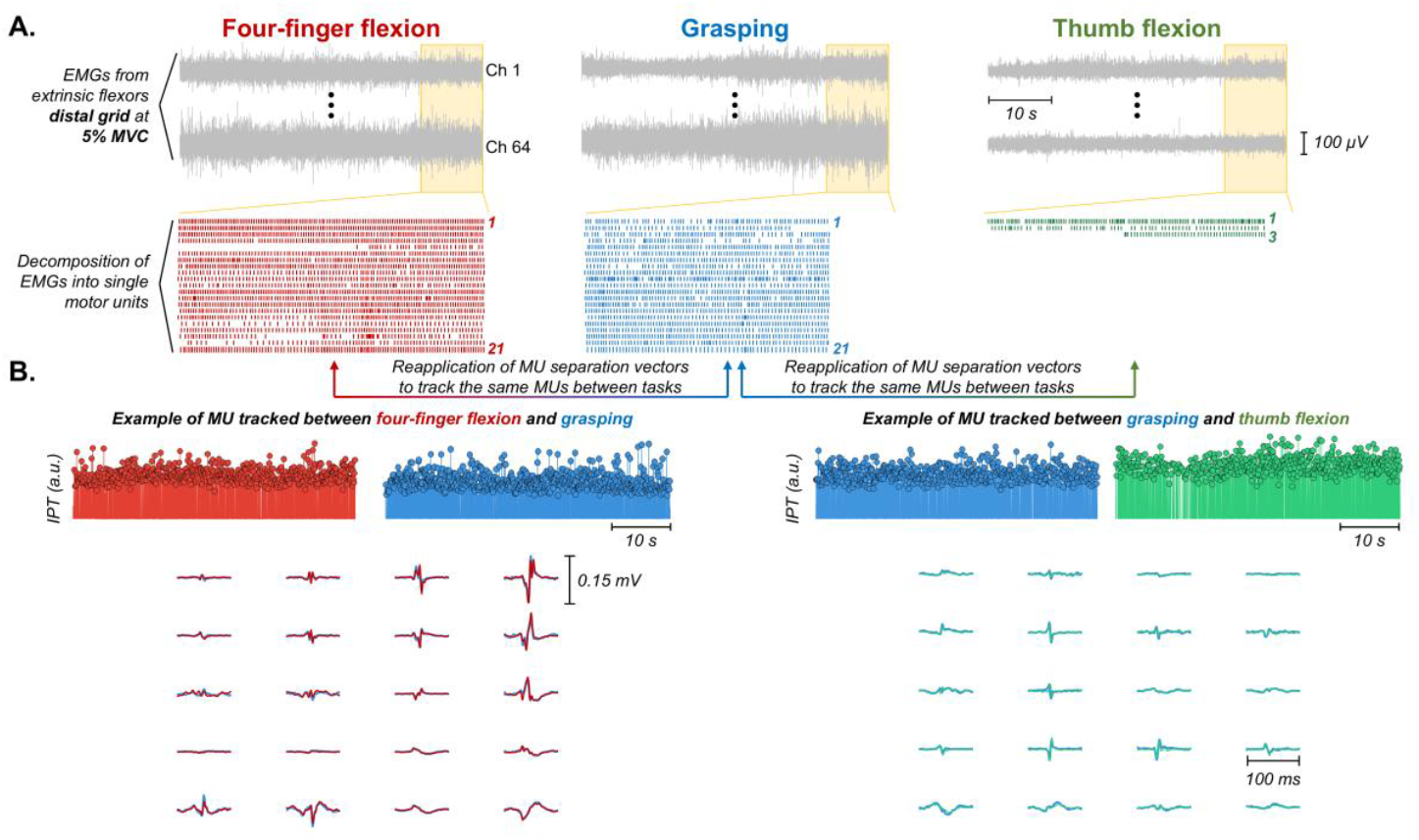
Data analysis. A, high-density surface electromyograms acquired during four-finger flexion (red), thumb flexion (green), and grasping (blue) were decomposed into motor unit spike trains using a convolutive blind source separation algorithm (Negro et al., 2016). Raster plots of decomposed motor units for each task are shown. Note that, for visualization purposes, the raster plots are shown for the last 10 s of the task (yellow boxes). B, motor units were tracked between tasks by reapplying the motor unit separation vectors from one contraction to the other, and vice-versa, so the same motor units were analyzed across four-finger flexion and grasping, as well as between thumb flexion and grasping. Examples of the innervation pulse trains (i.e., motor unit discharge times) and the two-dimensional representations on twenty selected single-differential channels of the motor units tracked between tasks are shown.

#### Estimates of common synaptic oscillations

Coherence analysis between motor unit spike trains was used to estimate common synaptic oscillations to spinal motor neurons (Negro and Farina, 2012, Castronovo et al., 2015, Cabral et al., 2024a, Cabral et al., 2024b). Due to the low number of matched motor units between thumb flexion and grasping tasks (see *Results*), only matched motor units between four-finger flexion and grasping were included in this analysis. Specifically, two equally-sized cumulative spike trains (CSTs), obtained as the sum of binary discharge times from randomly selected motor units, were used to calculate the coherence. Each CST comprised half of the total number of matched motor units, with this process repeated up to 100 random permutations (Cabral et al., 2024b). Only participants with at least four matched motor units were included in the coherence analysis (Cabral et al., 2024a). For each interaction, coherence was estimated between the two detrended CSTs using Welch’s periodogram with 95% overlapping Hanning windows of 1 s. This analysis was performed over a 30-s time window during the steady contraction, which was selected to maximize the number of active motor units (Maillet et al., 2022).

The obtained coherence values were averaged across interactions (i.e., pooled coherence) and z-transformed as previously described (Gallet and Julien, 2011). Only values higher than the bias, defined as the mean value of z-scores between 250 and 500 Hz (Castronovo et al., 2015, Maillet et al., 2022), were considered for further analysis. To assess changes between four-finger flexion and grasping tasks, the area under the curve of the z-coherence was calculated separately for the delta (1-5 Hz), alpha (5-15 Hz) and beta (15-35 Hz) bands. Then, the area under the curve ratio (grasping/four-finger flexion) was computed and 1 was subtracted (Cabral et al., 2024a). Thus, ratio values higher than and lower than 0 indicated, respectively, increases and decreases in z-coherence during grasping compared with the four-finger flexion task.

#### Coupling between neural drive and force oscillations

Force signals were initially low-pass filtered using a 3^rd^ order Butterworth filter with a cutoff frequency of 15 Hz. To assess potential changes in the coupling between neural drive and force oscillations when comparing four-finger flexion and grasping tasks, coherence was calculated between the neural drive and force signals (Negro et al., 2009, Cabral et al., 2024b). The neural drive to the superficial extrinsic flexor muscles of the hand was estimated as the CST of all identified motor units (Thompson et al., 2018). Similar to the analysis of common synaptic inputs, the area under the curve ratio (grasping/four-finger flexion) of z-coherence was calculated to evaluate changes between four-finger flexion and grasping tasks, but focusing only on the frequency bandwidths relevant for force production (i.e., delta and alpha bands; (Cabral et al., 2024b)). Additionally, to quantify alterations in the force steadiness between tasks, we measured the coefficient of variation of force.

### Statistical analysis

All statistical analyses were conducted in R (version 4.3.0) within the RStudio environment (version 2023.03.1).

Linear mixed-effect models (LMM) were used to compare the mean discharge rate between four-finger or thumb flexion and grasping tasks, separately for each force level (5% and 15% MVC). This statistical approach accounts for the hierarchical nature of motor unit data, recognizing the non-independence of observations (Tenan et al., 2014), and allows for the inclusion of all matched motor units rather than a single mean value for each participant. Random intercept models were used with ‘task’ as the fixed effect and ‘participant’ as the random effect. LMMs were implemented using the *lmerTest* package (Kuznetsova et al., 2017) with the Kenward-Roger method employed to approximate the degrees of freedom and estimate the *P*-values. The *emmeans* package was applied to determine estimated marginal means and their differences with 95% confidence intervals (Lenth et al., 2019).

To compare estimates of common synaptic inputs (i.e., z-coherence between motor unit spike trains) and CST-force coupling between four-finger flexion and grasping tasks, changes in the area under the curve ratio (grasping/four-finger flexion) were statistically tested using one-sample Wilcoxon signed-rank test (null hypothesis; μ0 = 0), separately for each frequency bandwidth. Additionally, to compare the coefficient of variation of force between four-finger flexion and grasping tasks, Wilcoxon signed-rank tests were used separately for each force level.

The effect sizes, along with their 95% confidence intervals, are reported for all the results. For the results of motor unit mean discharge rate, effect sizes were calculated using the *eff_size* function in R, which estimates Cohen’s d directly from the outputs of the *emmeans* package. For the area under the curve ratio results, effect sizes were calculated using Cohen’s d_s_ as reported in Jané et al. (2024). For the coefficient of variation of force results, effect sizes were calculated using Cohen’s d_z_ as reported in Jané et al. (2024).

For the results of motor unit mean discharge rate, the values in the text are reported as mean with 95% confidence intervals. All the other values are reported as median/interquartile ranges unless indicated otherwise. All individual data on matched motor unit discharge times are available at https://doi.org/10.6084/m9.figshare.28923704.

## Results

### Motor unit identification and mean discharge rate

To compare changes in the mean discharge rate of motor units between four-finger or thumb flexion and grasping tasks, we decomposed HDsEMG from the superficial extrinsic flexor muscles of the hand. For 5% MVC, 20 ± 8 motor units were identified per participant during four-finger flexion, 12 ± 9 during thumb flexion, and 19 ± 7 during the grasping task. For 15% MVC, 22 ± 8 motor units per participant during four-finger flexion, 12 ± 9 during thumb flexion, and 22 ± 8 during grasping were identified. Then, the identified motor units between tasks were tracked to ensure the same units were analyzed. For 5% MVC, 9 ± 6 units were matched between four-finger flexion and grasping tasks, and 2 ± 2 units between thumb flexion and grasping tasks. For 15% MVC, these numbers were 10 ± 8 between four-finger flexion and grasping, and 2 ± 4 between thumb flexion and grasping.

Significant reductions in motor unit mean discharge rate were observed between four-finger flexion and grasping tasks for both force levels. For 5% MVC, the mean discharge rate decreased from 12.7 [11.8, 13.5] to 12 [11.1, 12.8] pps (LMM, F = 9.15, *P* = 0.003, Cohen’s d = -0.35 [-0.59, -0.11]), and for 15% MVC, from 14.5 [13.4, 15.5] to 13.6 [12.5, 14.6] pps (LMM, F = 18.64, *P* < 0.001, Cohen’s d = -0.47 [-0.72, -0.24]). Conversely, the mean discharge rate significantly increased between thumb flexion and grasping tasks for both 5% MVC (from 11.5 [10.3, 12.7] to 13.2 [12, 14.4] pps; LMM, F = 14.38, *P* < 0.001, Cohen’s d = 0.89 [0.36, 1.43]) and 15% MVC (from 12.6 [11.6, 13.6] to 13.6 [12.5, 14.6] pps; LMM, F = 4.16, *P* = 0.045, Cohen’s d = 0.48 [0.04, 1]) force levels.

### Estimates of common synaptic inputs

To investigate changes in common synaptic inputs, we calculated the z-coherence between motor unit spike trains. The ratio of the area under the curve was then quantified separately for the delta (1-5 Hz), alpha (5-15 Hz) and beta (15-35 Hz) bands. For this analysis, only matched motor units between four-finger flexion and grasping tasks were utilized, as very few units were matched between thumb flexion and grasping. Additionally, only participants with at least four matched motor units were included. Consequently, coherence analysis was performed for 12 and 11 participants for 5% and 15% MVC, respectively. **Figure 4** displays the pooled z-coherence for all participants and the area under the curve ratio results. For both 5% MVC (top panel) and 15% MVC (bottom panel) force levels, there was a clear decrease in the area under the curve in the alpha band (yellow area) for the grasping task (blue) compared to the four-finger flexion task (red). For 5% MVC, a significant median decrease of ∼15% was observed in the area under the curve within the alpha band (**Figure 3A**; one-sample Wilcoxon signed-rank test, *P* = 0.009, Cohen’s d_s_ = -0.81 [-1.45, -0.14]), but no significant changes were found for the delta (one-sample Wilcoxon signed-rank test, *P* = 0.622, Cohen’s d_s_ = 0.20 [-0.38, 0.77]) or beta bands (one-sample Wilcoxon signed-rank test, *P* = 1.000, Cohen’s d_s_ = 0.01 [-0.56, 0.58]). Similarly, for 15% MVC, the area under the curve within the alpha band for the grasping task was significantly decreased compared to the four-finger flexion task by a median of ∼22% (**Figure 3B**; one-sample Wilcoxon signed-rank test, *P* = 0.032, Cohen’s d_s_ = -0.87 [-1.56, - 0.16]), with no significant changes in the delta (one-sample Wilcoxon signed-rank test, *P* = 0.700, Cohen’s d_s_ = 0.20 [-0.41, 0.79]) or beta bands (one-sample Wilcoxon signed-rank test, *P* = 0.700, Cohen’s d_s_ = 0.08 [-0.52, 0.67]).

**Figure 3:**
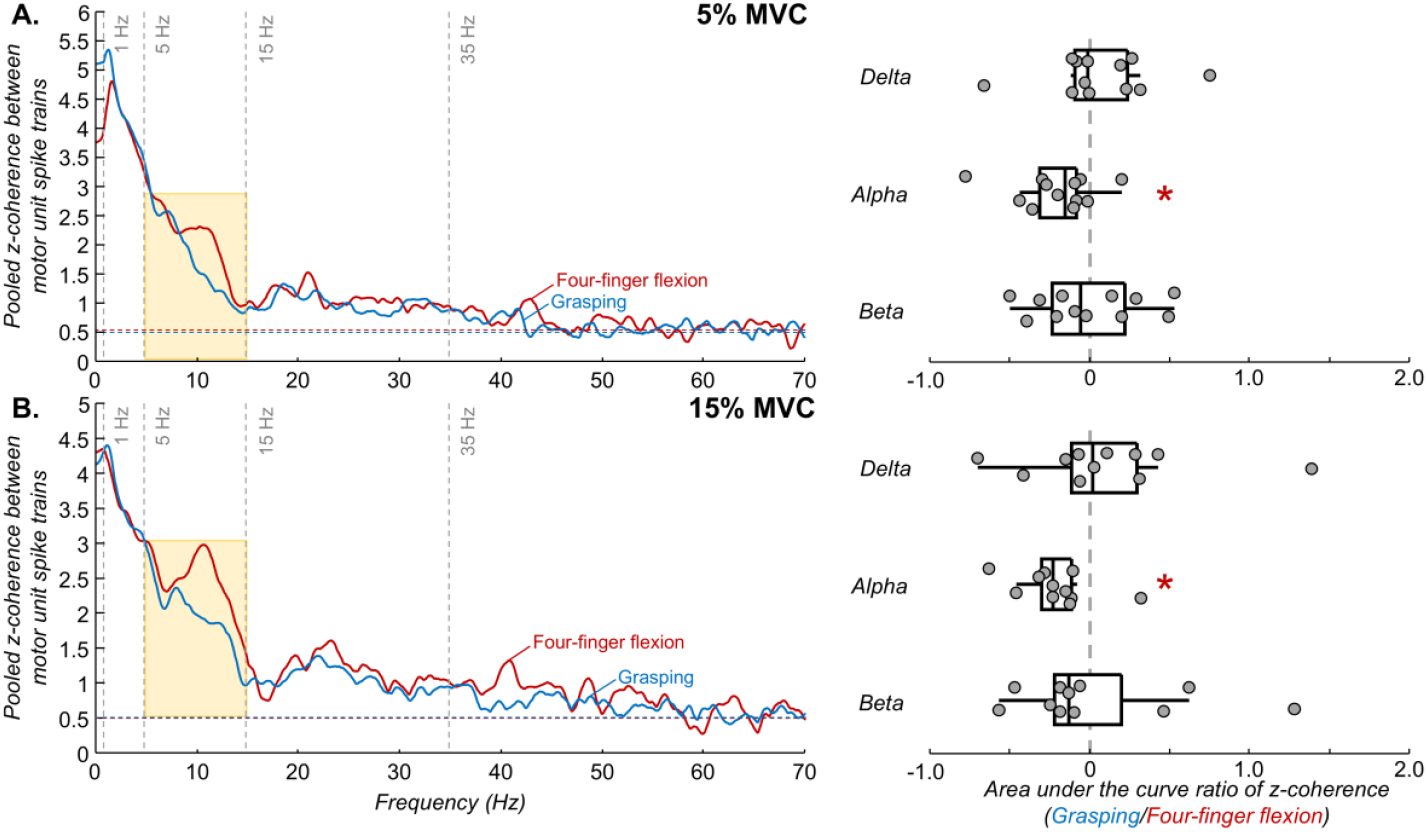
Motor unit coherence results. The left panel shows pooled z-coherence profiles including all participants for 5% MVC (A) and 15% MVC (B). Red and blue traces denote four-finger flexion and grasping tasks, respectively. The horizontal dashed line indicates the confidence level of coherence. Vertical dashed lines highlight the three frequency bandwidths analyzed: delta (1–5 Hz), alpha (5–15 Hz), and beta (15–35 Hz) bands. Yellow boxes denote statistical differences in the area under the curve between four-finger flexion and grasping tasks. The right panel displays group results of the area under the curve ratio of coherence for 5% MVC (top) and 15% MVC (bottom). Circles identify individual participants. Vertical black traces, boxes, and whiskers denote the median value, interquartile interval, and distribution range, respectively. *P < 0.05.

**Figure 4:**
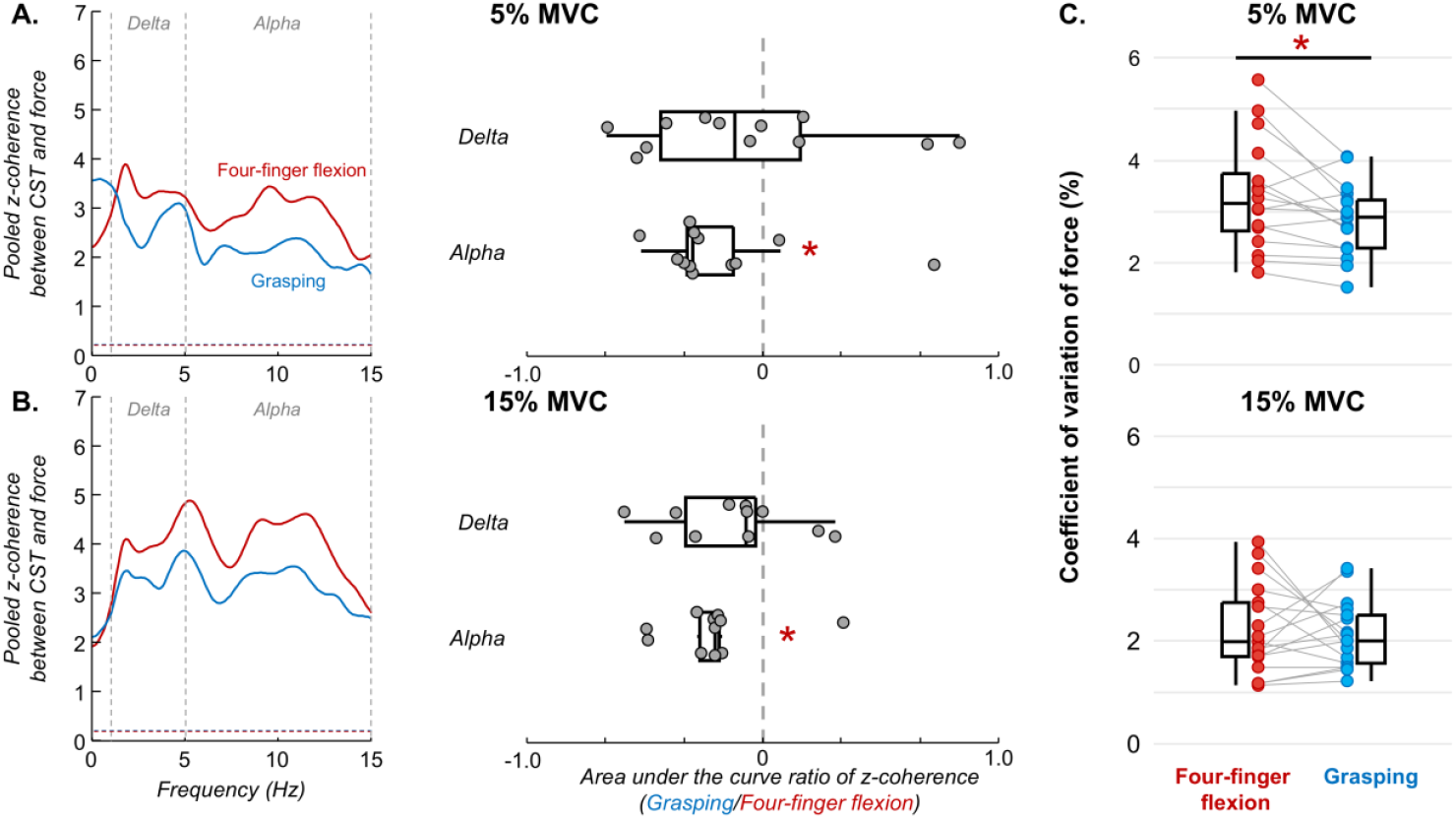
Results of force steadiness and coherence between neural drive and force oscillations. A-B, left panel shows pooled z-coherence profiles between the motor units cumulative spike train (CST) and force for 5% MVC (A) and 15% MVC (B). Red and blue traces denote the four-finger flexion and grasping tasks, respectively. The horizontal dashed line indicates the confidence level of coherence. Vertical dashed lines highlight the two frequency bandwidths analyzed: delta (1–5 Hz) and alpha (5–15 Hz) bands. Note that the frequency bandwidth analyzed was limited to the frequency bandwidth of force. The right panel shows group results of the area under the curve ratio of z-coherence for 5% MVC (top) and 15% MVC (bottom). Circles identify individual participants. Vertical black traces, boxes, and whiskers denote the median value, interquartile interval, and distribution range, respectively. C, group results of the coefficient of variation of force (force steadiness) for 5% MVC (top) and 15% MVC (bottom). Coloured circles and horizontal grey lines indicate individual participants. *P < 0.05.

### Coupling between neural drive and force oscillations

To determine if the observed changes in the alpha band of common synaptic inputs between the four-finger flexion and grasping tasks were accompanied by alterations in the linear coupling between the neural drive and the force oscillations, we calculated the z-coherence between the CST and the force signals. For this analysis, only the bandwidths relevant to force production were examined (delta and alpha bands). For both 5% MVC (**Figure 4A**) and 15% MVC (**Figure 4B**) force levels, significant decreases in the CST-force z-coherence during the grasping task were observed within the alpha band (one-sample Wilcoxon signed-rank test; 5% MVC: *P* = 0.043, Cohen’s d_s_ = -0.55 [-1.15, -0.07]; 15% MVC: *P* = 0.032, Cohen’s d_s_ = -0.97 [-1.68, -0.23]), with no significant changes for the delta band (one-sample Wilcoxon signed-rank test; 5% MVC: *P* = 0.519, Cohen’s d_s_ = 0.20 [-0.41, 0.79]; 15% MVC: *P* = 0.102, Cohen’s d_s_ = -0.49 [-1.11, 0.15]).

To assess changes in force steadiness between four-finger flexion and grasping tasks, we calculated the coefficient of variation of force. Significant reductions in the coefficient of variation of force were observed between the four-finger flexion and grasping tasks for 5% MVC (**Figure 4C**; Wilcoxon signed-rank test, *P* = 0.009, Cohen’s d_z_ = -0.41 [-0.90, 0.09]), but not for 15% MVC; Wilcoxon signed-rank test, *P* = 0.854, Cohen’s d_z_ = -0.11 [-0.58, 0.37]) force level.

## Discussion

The main purpose of this study was to investigate changes in common synaptic inputs to spinal motor neurons between a functionally relevant and frequently used grip (grasping) and a less functionally relevant hand task (four-finger flexion without thumb involvement). Our results demonstrated a reduction in alpha band (5–15 Hz) oscillations during grasping compared to the four-finger flexion task. These differences were reflected in force oscillations, revealing decreased force-neural drive coupling within the alpha band and increased force steadiness during grasping. These findings suggest that a more functionally relevant and natural grip involves specific neural control mechanisms to enhance force control.

Considering the complex biomechanical and neural structure of the hand, even frequently performed daily tasks, such as grasping an object, require precise coordination of multiple degrees of freedom that must be controlled simultaneously (Fuglevand, 2011, Santello et al., 2013). Over the past three decades, researchers have sought to understand how the fine coordination of multiple hand muscles is achieved during finger and hand movements (Schieber, 1990, Kilbreath and Gandevia, 1994, Santello and Soechting, 2000, Keen and Fuglevand, 2004, Hockensmith et al., 2005, Tanzarella et al., 2021). As reviewed by Schieber (1990), two (non-mutually exclusive) models describe how descending pathways may be organized to achieve this control. In the first model, each muscle or digit is controlled individually, with corticospinal pathways projecting separately to each motor neuron pool. In the second model (proposed as model 3 in Schieber (1990)), a small set of muscle activity patterns, or synergies, generate the full repertoire of hand movements, with corticospinal pathways diverging across individual motor neuron pools within and across muscles. Compelling evidence from force (Li et al., 1998, Reilly and Hammond, 2000, Santello and Soechting, 2000), kinematic (Soechting and Flanders, 1997, Santello et al., 1998, Santello et al., 2002), muscle activation (Maier and Hepp-Reymond, 1995, Weiss and Flanders, 2004, Tanzarella et al., 2021), and motor unit (Hockensmith et al., 2005, Tanzarella et al., 2021, Winges and Santello, 2004) recordings suggest that the latter model likely exemplifies hand control, with the synaptic inputs that are shared across spinal motor neurons playing a key functional role in grip force coordination (Santello and Fuglevand, 2004). However, the degree of commonality across motor units within and between hand muscles appears to be muscle- and task-dependent. For instance, while extrinsic hand muscles show a significant degree of common synaptic inputs between motor units, some suggest that intrinsic hand muscles exhibit lower commonality (Winges and Santello, 2004, Hockensmith et al., 2005, McIsaac and Fuglevand, 2008, Gandevia and Rothwell, 1987). Interestingly, in the current study it was demonstrated that grip type – highly functional (grasping) versus less frequently performed (four-finger flexion) – also modulates common synaptic inputs across motor units of the superficial extrinsic flexor muscles of the hand. Specifically, we observed a reduction in alpha band coherent oscillations between tracked motor units during grasping compared to four-finger flexion (**Figure 3**).

Although the physiological basis of the reduction in alpha band oscillations during grasping remains challenging to pinpoint, several potential mechanisms could explain this finding. First, the addition of thumb flexion alongside four-finger flexion (i.e., grasping) compared to four-finger flexion alone may introduce an additional source of common synaptic input to spinal motor neurons, leading to a reduction in alpha band coupling. This hypothesis aligns with previous evidence that reported when a new common input is introduced independently to a motor neuron pool, it can decorrelate motor unit spike train output within specific frequency bands (Negro and Farina, 2011). Second, previous studies suggested that spinal interneurons may phase-cancel cortical oscillatory inputs in the alpha frequency range, thereby reducing force tremor, which subsequently enhances movement precision (Williams et al., 2010, Koželj and Baker, 2014, Cabral et al., 2024a). This mechanism could contribute to improved force control in grasping, which is a more frequently used movement compared to four-finger flexion. Finally, given that modulations in the gain of the Ia afferent feedback loop can directly influence alpha band inputs (Halliday and Redfearn, 1956, Lippold, 1971, Christakos et al., 2006, Laine et al., 2016), it is possible that grasping involves a reduction in Ia afferent gain. Since grasping is more frequently performed in daily life, it may rely less on afferent feedback compared to a less applied and less functional grip. Nevertheless, it remains possible that more complex neural mechanisms are involved, integrating both cortical and spinal pathways.

Our findings have directly applied relevance, as the observed reductions in alpha band oscillations during grasping were reflected in force fluctuations. Since alpha band fluctuations in the effective neural drive to the muscle are not fully canceled by the muscle’s twitch contractile properties, which act as a low-pass filter with a cutoff frequency of ∼12 Hz (Bawa and Stein, 1976, Baldissera et al., 1998), they are consequently transmitted to force, contributing to its variability (Negro et al., 2009, Farina et al., 2014). Thus, the reduction in alpha band coherence during grasping may represent a neural strategy to enhance force steadiness in a frequently performed hand task, effectively filtering out frequency oscillations from the control signal responsible for force modulation (i.e., delta band coherence) (Farina et al., 2014, Hug et al., 2023). Supporting this interpretation, the observed decrease in the coefficient of variation of force (indicative of increased force steadiness) during grasping with respect to four-finger flexion at low force levels (**Figure 4C**). This was further supported by a decrease in alpha band coupling between the cumulative spike train and force during grasping compared to four-finger flexion (**Figure 4A** and **4B**). These results align with previous studies and support the findings that various factors, such as sensorimotor integration (Laine et al., 2014), muscle contractile properties (Cabral et al., 2024b), pain (Yavuz et al., 2015), or learning a new motor skill (Cabral et al., 2024a), modulate the alpha band in the common synaptic input to motor neurons, directly influencing optimal force control. Therefore, our findings extend this evidence by demonstrating that grip type affects common synaptic oscillations to spinal motor neurons, further influencing force regulation in functionally relevant tasks.

While these experiments do not allow us to directly address this question, a key issue arising from the findings in this study is whether the observed modulation in common synaptic inputs reflects a hard-wired neural control strategy or emerges due to task constraints (i.e., soft- assembled control), which is a topic of ongoing debate in the literature (Tresch and Jarc, 2009, Kutch and Valero-Cuevas, 2012, Dernoncourt et al., 2025). Recently, using a combination of kinematic, electromyography, and functional magnetic resonance imaging data, Leo et al. (2016) demonstrated that human motor cortical areas encode hand movement synergies, supporting the idea that modular hand control is represented at higher motor centers. Similarly, Takei et al. (2017) reported that modular control of hand movements is implemented at the spinal level in primates, suggesting that the nervous system organizes motor outputs into functional modules to simplify the control of voluntary hand movements. In addition to this evidence, given that grasping is an innate reflex in infants, which later develops into voluntary grasping (Dennis, 1943, Twitchell, 1965), it seems to follow that modulation of common synaptic oscillations observed in this study may be embedded in the neural circuitry. However, further research is needed to validate this hypothesis.

A final consideration regarding differences in motor unit mean discharge rates. This study reported a reduction in mean discharge rate during grasping compared to four-finger flexion alone, though with relatively small effect sizes (see Results). Given the complexity of intrinsic and extrinsic muscle coordination required for hand control, some motor units within the extrinsic flexor muscles primarily contribute to force production, while others play a role in force distribution across muscles and mechanical stabilization (Li, 2002, Kinoshita et al., 1995, Butler et al., 2005). Therefore, the observed small reduction in mean discharge rate may reflect mechanical adaptations rather than direct changes in neural control. Notably, this decrease was contrasted by an increase in mean discharge rate when the thumb was flexed alone compared to grasping. This suggests that the nervous system dynamically adjusts motor unit activity, whether for force generation or mechanical stabilization, to maintain the required force target across different grip conditions.

In conclusion, the findings of this study demonstrate that grasping, which is a functionally relevant and habitually performed hand movement, exhibits distinct neural control mechanisms compared to less commonly used grips. Therefore, the reduction in alpha band oscillations during grasping suggests a task-dependent modulation of common synaptic input, which has functional implications for enhanced force steadiness and optimized neural drive-force coupling.

## Acknowledgements

This study was funded by the European Research Council Consolidator Grant INcEPTION contract no. 101045605. J Greig Inglis was supported by the Marie Skłodowska-Curie Actions Grant ‘MUDecomp’, agreement no. 101151712.

